# Evaluation of KRAS^G12C^ Inhibitor Responses in Novel Murine KRAS^G12C^ Lung Cancer Cell Line Models

**DOI:** 10.1101/2022.08.19.504555

**Authors:** Daniel J Sisler, Trista K Hinz, Anh T Le, Emily K Kleczko, Raphael A Nemenoff, Lynn E Heasley

## Abstract

The KRAS(G12C) mutation is the most common genetic mutation in North American lung adenocarcinoma patients. Recently, direct inhibitors of the KRAS^G12C^ protein have been developed and demonstrate clinical response rates of 37-43%. Importantly, these agents fail to generate durable therapeutic responses with median progression-free survival of ~6.5 months. To provide models for further preclinical improvement of these inhibitors, we generated three novel murine KRAS^G12C^-driven lung cancer cell lines. The co-occurring NRAS^Q61L^ mutation in KRAS^G12C^-positive LLC cells was deleted and the KRAS^G12V^ allele in CMT167 cells was edited to KRAS^G12C^ with CRISPR/Cas9 methods. Also, a novel murine KRAS^G12C^ line, mKRC.1, was established from a tumor generated in a genetically-engineered mouse model. The three lines exhibit similar *in vitro* sensitivities to KRAS^G12C^ inhibitors (MRTX-1257, AMG-510), but distinct *in vivo* responses to MRTX-849 ranging from progressive growth with orthotopic LLC-NRAS KO tumors to marked shrinkage with mKRC.1 tumors. All three cell lines exhibited synergistic *in vitro* growth inhibition with MRTX-1257 and the SHP2 inhibitor, RMC-4550 and the MRTX-849/RMC-4550 combination yielded tumor shrinkage in orthotopic LLC-NRAS KO tumors propagated in syngeneic mice. Notably, this synergistic combination response was lost in athymic *nu/nu* mice, supporting a growing literature demonstrating a role for adaptive immunity in the response to this class of drugs. These new models of murine KRAS^G12C^ mutant lung cancer should prove valuable for identifying improved therapeutic combination strategies with KRAS^G12C^ inhibitors.

**Contribution to the Field Statement:** The development of KRAS^G12C^ inhibitors has not impacted treatment of lung cancers bearing the KRAS^G12C^ mutation to the degree that tyrosine kinase inhibitors have changed the treatment outcomes for patients bearing oncogenic mutations in receptor tyrosine kinases. Thus, the field is now exploring combination strategies with KRAS^G12C^ inhibitors that may enhance their clinical benefit. Moreover, published findings indicate that host immunity contributes to efficacy of oncogene-directed inhibitors including KRAS^G12C^ inhibitors. Thus, these novel murine KRAS^G12C^-driven lung cancer cell lines will provide valuable models for preclinical evaluation of novel drug combinations in immune competent hosts.

## Introduction

Lung cancer is the second most common cancer diagnosed in men and women, and accounts for the highest proportion of cancer-related deaths^1^. In 2022, an estimated 240,000 patients will be diagnosed with lung cancer, resulting in an estimated 130,000 deaths^1^. Lung adenocarcinomas (LUAD) account for ~ 30-40% of lung cancer patients and are clinically defined by their oncogenic driving mutations^2^. Recent advances in precision medicine have led to the clinical approval of small molecule inhibitors that directly target oncogenic forms of EGFR, ALK, ROS1, RET, NTRK and BRAF mutant cancers. While pharmacological targeting of oncogenic receptor tyrosine kinases (RTKs) has experienced great success, other dominant oncogenic drivers including KRAS could not be directly targeted until recently. Mutations in the KRAS oncogene occur in ~30% of LUAD and inhibitors targeting the most common mutational subset of this gene (glycine to cysteine at codon 12) have been developed and gained FDA approval for use in a second line setting^3,4^. Early clinical data demonstrate that the KRAS^G12C^ inhibitor sotorasib (AMG-510) achieved an objective response rate of 37.1% and the median duration of response was 11.1 months with a comparatively short progression free survival of 6.8 months^5^. These data indicate that single agent KRAS^G12C^ inhibitor activity is short-lived and as previous research has shown, ERK reactivation through both bypass signaling and acquired resistance mechanisms may drive these abbreviated responses^6,7^. These findings argue for the development of rational therapeutic combinations that may prolong ERK inhibition in a targeted manner to bolster duration of response.

KRAS^G12C^ inhibitors function to covalently bind KRAS^G12C^ in its GDP-bound (inactive) state. Distinct from other KRAS codon 12 alleles, KRAS^G12C^ retains intrinsic GTP hydrolysis capabilities^8^, highlighting the targeting of proteins upstream of KRAS signaling as attractive candidates to target in combination with KRAS^G12C^ inhibitors^6^. This upstream targeting helps prevent KRAS activation and nucleotide exchange for GTP, thereby making more GDP-bound KRAS available for KRAS^G12C^ inhibitor to bind. The tyrosine-protein phosphatase SHP2 has emerged as a promising therapeutic target^9^. This protein acts as a mediator associated with the downstream stimulation of the RAS/RAF/MEK/ERK pathway that promotes MAPK signal activation by dephosphorylating activating phospho-tyrosine residues within the RAS-GAPs, NF1 and p120RASGAP^10–12^. SHP2 inhibition alone has little effect on reducing KRAS-driven tumor cell growth, but in combination with a MEK inhibitor resulted in synergistic tumor shrinkage^13^. Another attractive aspect of targeting SHP2 is its functional repression of JAK/STAT and immune related signaling pathways^14^. Targeting this protein with allosteric SHP2 inhibitors promotes anti-tumor immunity, including enhancement of T cell cytotoxic function and immune-mediated tumor regression via a variety of different mechanisms^14^. SHP2 has also been shown to abrogate KRAS^G12C^ inhibitor-specific responses in some models^15^. Clinically, SHP2 inhibitors have exhibited encouraging disease control rates of 67% for advanced NSCLC patient with KRAS mutations, although significant adverse events related to toxicity were observed. Still, the ability to target both mutant KRAS signaling, as well as immune related signaling makes this combination appealing.

Our lab and others have shown the importance of innate and adaptive immunity in driving pre-clinical and clinical responses to oncogene-targeted therapies^16–22^. Thus, comprehensive assessment of the therapeutic potential of oncogene-targeted agents requires immune competent murine models. To date, few murine models of KRAS^G12C^ lung cancer exist to test the effects of KRAS^G12C^ inhibitor in immune competent hosts. Herein, we have developed three new murine KRAS^G12C^-dependent lung cancer cell lines and tested their sensitivity to a KRAS^G12C^ inhibitor as a monotherapy and in combination with a SHP2 inhibitor *in vitro* as well as *in vivo* using an orthotopic model of lung tumor growth. The findings reveal superior activity of the drug combination in immune competent, but not immune-deficient mice.

## Materials and Methods

### Cell Culture

All human and murine cell lines were cultured in RPMI-1640 media (Corning, Tewksbury, MA) supplemented with 5% fetal bovine serum (FBS) and 1% penicillin–streptomycin (Sigma-Aldrich, St. Louis, MO) at 37 °C in a humidified 5% CO_2_ incubator. All human cell lines used in this study were submitted to fingerprint analysis to confirm authenticity within a year of performing the studies described herein. A reference STR genotype was acquired through IDEXX Bioresearch for all murine lines and was later used to confirm genotypic integrity. CMT167 cells^23^ and Lewis Lung Carcinoma cells (LLC) were obtained from the lab of Dr. Raphael Nemenoff. All cell lines were periodically tested for mycoplasma infection. To avoid cross-contamination and phenotypic changes, cells were maintained as frozen stocks and cultured for only two to four weeks before use in experiments. Authentication of cell lines based on morphology and growth curve analysis was performed regularly. No phenotypic changes were observed through the duration of the study.

### KRAS^G12C^ Mouse Tumor Cell Line Generation

B6;129S4-*Kras*^*em1Ldow*^/J mice (Lukas Dow, Jackson Labs) were crossed with B6.129P2-*Trp53*^*tm1Brn*^/J mice. F1 hybrid mice were genotyped for KRAS and p53 status using PCR on genomic DNA and the following primer pairs: KRAS^G12C^, LSL-Kras Common 5’-TCCAATTCAGTGACTACAGATG-3’, LSL-Kras Mutant 5’-CTAGCCACCATGGCTTGAGT-3’, LSL-Kras Wildtype 5’-ATGTCTTTCCCCAGCACAGT-3’); Trp53, FWD 5’-GGTTAAACCCAGCTTGACCA-3’, REV-5’-GGAGGCAGAGACAGTTGGAG-3’). All PCR reactions employed PCRBIO VeriFi™ mix (PCRBIO, Wayne, Pennsylvania) according to manufacturer’s protocol with 60°C annealing temperature. Mice heterozygous for KRAS^G12C^ and varying p53 backgrounds (p53^+/+^, p53^+/flox^, and p53^flox/flox^) were inoculated with Ad-Cre intratracheally and submitted to microCT imaging once monthly until tumor burden was detected. Tumors were harvested, minced, plated on tissue culture plastic and cultured until the stable cell line, mKRC.1, was obtained.

### CRISPR-Cas9 Genome Editing

#### NRAS knockout

A crRNA was designed targeting exon 5 of genomic mouse NRAS using IDT (IDT, Iowa, City, IA) online software (5’-CACGAACTGGCCAAGAGTTA-3’). LLC cells (30,000) were plated in a 35 mm^2^ plate and cultured for 16 hrs. The NRAS targeting crRNA, ATTO550-labelled tracrRNA and recombinant Cas9 were assembled into ribonucleoprotein (RNP) particles according to IDT manufacturer’s protocol. Assembled RNPs were packaged using Lipofectamine RNAiMAX transfection reagent (ThermoFischer Scientific) according to manufacturer’s instructions, and transfection mixture was added to LLC cells. After a 24-hour incubation, cells were trypsinized, rinsed with 5 mL of PBS, and resuspended into a single-cell suspension in 1mL of PBS. Single cells were flow sorted into single wells of 96-well plates using the MoFlo XDP100 cell sorter for ATTO550 positivity (Beckman Coulter) (University of Colorado Anschutz Medical Campus Cancer Center Flow Cytometry Core). For proper compensation of flow cytometry channels, unstained cell lines were used. Single cell clones were expanded, genomic DNA collected, and assessed for NRAS perturbations via PCR and Sanger Sequencing. Two independent clones were identified and indicated as LLC 23 NRAS KO and LLC 46 NRAS KO.

#### KRAS G12V to G12C

A crRNA targeting exon 1 of genomic mouse KRAS (5’-GACTGAGTATAAACTTGTGG-3’) using IDT online software was designed in addition to single stranded donor oligonucleotides (ssODN) homology-directed repair templates for both the positive and negative strands, the sequence for which is: 5’-ATGACTGAGTATAAACTTGTCGTCGTTGGAGCTTGCGGCGTAGGCAAGAGCGCCTTGACGATACAGCTAATTCAGA-3’. CMT167 KRAS^G12C^ cell lines were subject to the same protocol used for generating LLC NRAS knockout cell lines with the addition of two steps. 1. Prior to transfection, CMT167 cells were incubated with 30 µM HDR Enhancer (IDT) for 1 hour. 2. ssODN’s were added to the transfection mixture and co-transfected into cells along with assembled RNP’s. Single cell subclones from flow sorting were expanded and analyzed for endogenous KRAS^G12C^ mutations.

### RNA-seq and Bioinformatic Analysis

RNA was submitted to the University of Colorado Sequencing Core where library preparations were generated, and RNA was sequenced on the NovaSeq 4000 to generate 2 × 151 reads. Fastq files were quality checked with FastQC, illumina adapters trimmed with bbduk, and mapped to the mouse mm10 genome with STAR aligner. Counts were generated by STAR’s internal counter and reads were normalized to counts per million (CPM) using the edgeR R package. Differential expression was calculated using the limma R package and the voom() function. Heatmaps were generated in Prism 9 (GraphPad Prism Software, San Diego, CA). Integrated Genomics Viewer analysis was completed by stripping section of sequence reads from RNAseq dataset and viewing in IGV software^24^.

### Variant Calling

Fastq files were quality checked with FastQC, illumina adapters trimmed with bbduk, and paired reads were mapped to the UCSC mm10 BWA genome with samtools bwa-mem. Output sam files were converted to bam format, sorted, and duplicates were marked with the picard tool, MarkDuplicates. Samtools “mpileup” and bcftools “call” multiallelic caller were used to call variants and generate variant call files (VCF). To identify variants unique to each cell line, bcftools “isec” was used on each cell lines VCF files filtered with bcftools “view” excluding “G/T = 0/0” and including variants with at least a read depth of 10 and a quality score of 20 compared to unfiltered parental VCF files. Variants private to each cell line (001.vcf) were then annotated with ANNOVAR mm10db including refGene, cytoBand, and genomicSuperDups. VCFs were filtered excluding identified “intronic”, and “intergenic” annotated variants under the “Func.refGene tab in addition to “synonymous”, “.”, and “other” under the “ExonicFunc.refGene” tab. Interesting variants were confirmed by pulling out specific regions from the BAM files, uploading those reads to the integrated genomics viewer (IGV). A variant was considered detected reliably if the variant was present in > 10% of the total read depth at the variant position. All code and the parameters used are not included in this report but can be provided upon request.

### Cell Proliferation Assay

Cells were plated at 100 cells per well in 100 µL in 96 well tissue culture plates and allowed to attach for 24 hrs. Inhibitors were added at various doses as 2X concentrates in 100 µL. After incubation for 7-10 days, cell number per well was assessed using a CyQUANT Direct Cell Proliferation Assay (Invitrogen) according to the manufacturer’s instructions.

### Enzyme Linked Immunosorbent Assay (ELISA)

Conditioned media was collected from treated and untreated murine lung cancer cell lines. Chemokine levels were measured using the Invitrogen ELISA Kit (Quantikine mouse/human CXCL10/IP-10; R&D Systems, Minneapolis, MN) following manufacturer’s instructions.

Absorbance was measured at 450 nm. The measured concentration in each sample was normalized to the total cellular protein per dish and the data are presented as pg/µg protein.

### *In Vivo* Mouse Studies

Eight-week-old female C57B/6J mice were purchased from Jackson Labs (Bar Harbor, ME) and eight-week-old *nu/nu* mice were purchased from Envigo (Indianapolis, IN). For tumor cell inoculation, parental LLC, LLC 46 NRAS KO, and mKRC.1 cells were grown to 75% confluence, harvested, and resuspended in sterilized phosphate buffered saline. Cells were counted and resuspended to a final concentration of 125,000 cells per 40uL injection. Cell suspensions were directly injected into the left lung of mice through the ribcage. Mice were randomized into groups of 10 to receive diluent control, RMC-4550 (30 mg/kg; MedChem Express), MRTX-849 (30 mg/kg; MedChem Express), or the combination by daily oral gavage until primary experimental endpoints were met. Treatment was initiated following confirmation that at least 75% of mice had calculatable primary left lung tumors via microCT imaging (CUD Small Animal Imaging Core). MicroCT imaging was conducted weekly and ITK-SNAP software^25^ was used to calculate cross-sectional tumor volumes in cubic millimeters. Mice exhibiting signs of morbidity according to the guidelines set by the Institutional Animal Care and Use Committee (IACUC) were sacrificed immediately.

### Quantification and Statistical Analysis

Statistical Analysis: Prism 9 (GraphPad Software, San Diego, CA) was used to perform specific statistical analyses noted in the figure legends. Data are presented as the mean and standard error of the mean (SEM) as indicated. An unpaired Student’s t test (two-tail) was used to determine statistical significance, unless otherwise noted. The P values are denoted by *(PU<U0.05), **(PU<U0.01), ***(PU<U0.001), and ****(PU<U0.0001) and were corrected for multiple comparisons (Dunnett). Drug synergy with combinations of KRAS-G12C inhibitors and SHP2 inhibitors was determined with Combenefit, a free software tool for visualization, analysis and quantification of drug combination effects. Data from drug combination assays was processed using the HSA synergy model.

## Results

### Generation of murine lung cancer cell lines bearing KRAS^G12C^

The literature supports a role for innate and adaptive immunity in the overall tumoral responses to oncogene-specific inhibitors^16,21,26,27^. Herein, we generated three novel murine KRAS^G12C^-driven LUAD cell lines to permit investigation of KRAS^G12C^ inhibitor responses in immune-competent hosts.

#### LLC-NRAS-Q61L KO

Lewis lung carcinoma (LLC) are derived from a spontaneous lung tumor that arose in a C57BL/6 mouse^28^ and has been investigated by our group for sensitivity of orthotopic lung tumors to PD1/PD-L1 axis inhibitors^29–32^. Molina-Arcas et al. previously reported that a cell line derived from the LLCs (3LL) contained an NRAS-Q61H gain-of-function mutation in addition to a KRAS^G12C^ mutation^33^. Variant calling analysis (See Materials and Methods) of our published RNAseq data^29^ verified that our LLC cell line isolate also expressed NRAS-Q61L at the mRNA level (Supplementary Fig. S1A). To render this cell line fully dependent on KRAS^G12C^, a guide RNA (gRNA) targeting exon 5 of the murine NRAS gene was introduced into parental LLC as a complex with Cas9 and single-cell transfectants were isolated and screened for functional NRAS knockout. Genetic NRAS knockout (KO) in 2 independent LLC clones (LLC 23 NRAS KO and LLC 46 NRAS KO) was validated by RNAseq showing a large deletion near the gRNA targeting region of exon 5 of NRAS (Supplementary Fig. S1B).

#### CMT-167 KRAS-G12V to G12C

Like LLC cells, CMT-167 cells were isolated from a spontaneous lung tumor arising in a C57BL/6 mouse^23^ and bear an oncogenic KRAS-G12V mutation. CRISPR-Cas9-mediated, homology-directed repair was deployed to edit the endogenous G12V mutation to a G12C mutation. Ribonucleoprotein (RNP) particles containing Cas9, a single-stranded oligonucleotide DNA template and a gRNA targeting a sequence within exon 2 of murine KRAS were prepared and transfected into parental CMT cells. Following single cell flow sorting, clones were submitted to functional screening assessed for acquired sensitivity to KRAS^G12C^ inhibitors. Putative positive clones that successfully underwent homology-directed repair to contain the KRAS^G12C^ mutation were submitted to RNAseq and expression of the G12C allele was verified. RNA sequencing and IGV analysis confirmed an indel in one allele of murine KRAS and a successful recombination containing both KRAS^G12C^ as well as the predicted wobble position switches engineered to prevent Cas9 re-cutting of successfully recombined KRAS^G12C^ gene (Supplementary Fig. S1C).

#### Novel KRAS^G12C^ cell line from KRAS-G12C GEMM

KRAS^LSL-G12C/+^ mice developed by the Dow lab^34^ were obtained from Jackson Labs (B6;129S4-Kras^em1Ldow^/J) and crossed with Trp53^fl/fl^ mice (from Jackson Labs; B6.129P2-Trp53_tm1Brn_/J). The resulting KRAS^G12C/+^; Trp53^fl/fl^ mice were identified by genotyping and submitted to intratracheal administration of Ad-Cre as previously described^35^ (Supplementary Fig. S2A).

Approximately 29 weeks post Ad-Cre administration, dispersed and invasive solid lung tumors were identified in the dissected lungs of a mouse. The tumors were dispersed into single cells and passaged *in vitro* to yield the murine <u>KR</u>AS-G12<u>C</u> positive cell line, mKRC.1 (Supplementary Fig. S2A-B).

Variant calling of RNAseq data revealed 8 and 23 novel insertion/deletion mutations in LLC 23 NRAS KO and LLC 46 NRAS KO, respectively, that were not detected in any of the RNAseq reads from parental LLC cells (Supplemental Table S1). Analysis of the CRISPR/Cas9-edited CMT-167 lines revealed 25 and 36 novel insertion/deletion mutations in CMT KRAS-G12C.54.10 and CMT KRAS-G12C.55, respectively that were not detected in any of the RNAseq reads from parental CMT-167 cells (Supplementary Table S1). None of the indels reside in genes with reported functions in MAPK pathway signaling.

The LLC NRAS KO and CMT KRAS-G12C cell lines were submitted to *in vitro* proliferation assays to assess if the genetic perturbations altered baseline cell growth relative to the parental lines. A minimal effect of NRAS-Q61L knockout in LLC cells was observed on baseline proliferation rates (Supplementary Fig. S3A), although the rate of tumor growth of orthotopically implanted LLC 46 NRAS KO cells was markedly reduced relative to parental LLC cells (Supplementary Fig. S3B). Thus, the NRAS-Q61L significantly contributes to transformed growth properties of LLC cells. The edited CMT-KRAS-G12C cells exhibited significantly reduced *in vitro* proliferative rates compared to parental CMT-KRAS-G12V cells (Supplementary Fig. S3A) and is consistent with previous findings that KRAS^G12V^ has lower intrinsic GTP hydrolysis^8^ and cell lines expressing G12V mutations form more aggressive tumors than G12D or G12C mutations^36^. In addition, the CRISPR/Cas9-mediated editing of the CMT cells resulted in deletion of the unedited KRAS-G12V allele so that both clones bear a single copy of KRAS-G12C. Thus, reduced copy number of the oncogenic KRAS allele may contribute to the reduced basal growth rate. Pilot experiments with orthotopic inoculation of CMT-KRAS-G12C cells in syngeneic C57BL/6 mice revealed formation of very small primary tumors within the left lung lobe, but extensive growth in pleural and epicardial sites that required early euthanasia of the mice. As a result, *in vivo* studies with the CMT KRAS-G12C cell lines were not further pursued in this study.

### Sensitivity of novel KRAS^G12C^ murine cell lines to KRAS^G12C^ and MEK inhibitors

The sensitivity of the murine KRAS^G12C^ lung cancer cell lines to the KRAS^G12C^ inhibitors MRTX-1257 (Mirati Therapetics, San Diego, CA) and AMG-510 (Amgen, Thousand Oaks, CA) as well as the MEK1/2 inhibitor, trametinib was assessed with *in vitro* growth assays (see Materials and Methods). Knockout of NRAS increased the sensitivity of LLC NRAS KO cell lines to both KRAS^G12C^ inhibitors (AMG-510, 14-34 fold; MRTX-1257, 82-148 fold) compared to parental LLC cells (Fig. 1A), further supporting that mutated NRAS-Q61H significantly contributes as an oncogenic driver in these cells. This finding is consistent with that reported previously in the 3LL derivative of LLC cells^33^. LLC NRAS KO cell lines exhibited similar sensitivity to trametinib as parental LLCs (Fig. 1A), indicating that NRAS-Q61L serves as an additional proximal driver of MAPK signaling in LLC cells. Compared to parental CMT cells which are completely insensitive to MRTX-1257 and AMG-510 at concentrations of 3 nM and 300 nM, respectively, the CMT KRAS^G12C^-engineered cell lines were highly sensitive to these agents with IC_50_ values in the low nanomolar range (Fig. 1B). Interestingly, parental CMT cells bearing KRAS^G12V^ were less sensitive to trametinib than the CMT KRAS^G12C^ clones (Fig. 1B). This finding may reflect the disruption of the non-edited KRAS-G12V allele as a result of the CRISPR editing resulting in a single functional KRAS-G12C gene in the CMT-54 and CMT-55 clones. The reduced KRAS copy number may also contribute to their decreased proliferation rates observed *in vitro* (Suppl. Fig. S3A). Finally, the mKRC.1 cell line exhibits sensitivity to MRTX-1257 and AMG-510 (Fig. 1C). Analysis of MRTX-1257 and AMG-510-sensitivity in a panel of 13 human KRAS^G12C^-mutant lung cancer cell lines revealed IC_50_ values for these two drugs that ranged from 0.1 to 356 nM for MRTX-1257 and 0.3 to 2534 nM for AMG-510 (Supplementary Fig. S4A-B). The inhibitor sensitivity exhibited by the murine KRAS^G12C^ cell lines (Figure 1) overlapped with the most sensitive human lung cancer cell lines bearing KRAS^G12C^, indicating that co-occurring mutations present in many of the human lines markedly reduce their sensitivity to MRTX-1257 and AMG-510 similar to the activity of GTPase-deficient NRAS in parental LLC cells. Overall, the murine LUAD cell lines exhibit sensitivity to KRAS^G12C^ inhibitors that are consistent with their human counterparts.

**Figure 1.**
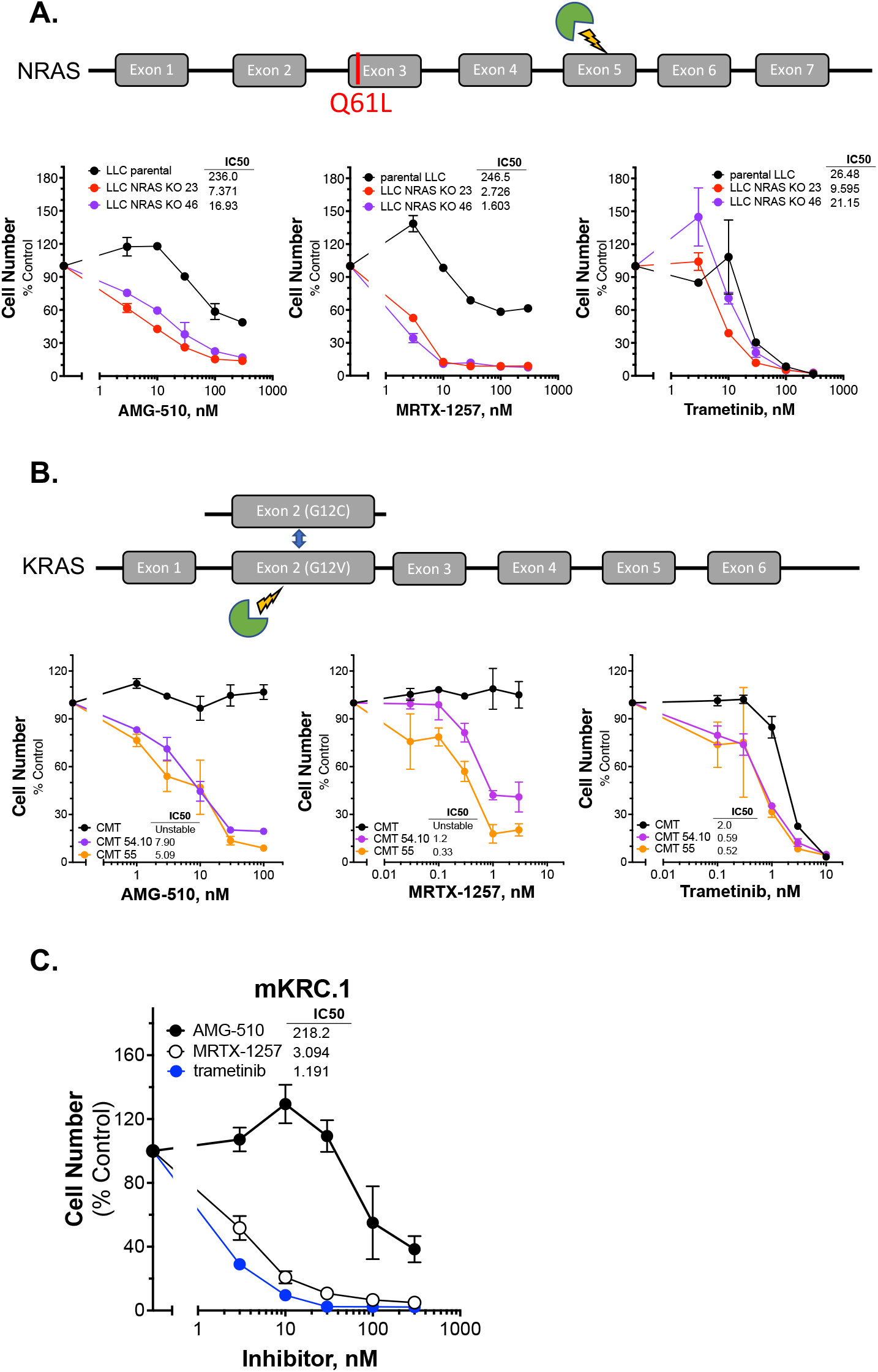
Murine KRAS G12C mutant lung cancer cell lines demonstrate sensitivity to KRAS and MEK inhibition. Schematic of CRISPR/Cas9 strategies are overlayed above each respective cell line panel. (**A**) Parental LLCs and 2 independent LLC clones harboring NRAS KO, (**B**) CMTs and 2 independent CMT clones with KRAS G12C substitutions. The parental and edited murine cell lines as well as (**C**) mKRC.1 were plated at 100-200 cells/well in triplicate in a 96-well plate, and then treated with increasing concentrations of the KRAS-G12C inhibitors MRTX-1257 and AMG-510, and the MEK inhibitor trametinib. Cell number was assessed 7-10 days later with the CyQuant assay. The data are the means ± SEM of triplicate determinations and presented as percent of the values measured in DMSO control wells. The experiments shown are representative of 2 to 3 independent experiments. The IC_50_ values were calculated with Prism 9 and presented within the individual dose-response curves as shown.

### SHP2 inhibition synergizes with KRAS^G12C^ inhibitors *in vitro* leading to enhanced chemokine response

In a pilot study, LLC 46 NRAS KO cells were propagated as orthotopic tumors in the left lung of C57BL/6 mice and treated with diluent or the MRTX-1257 clinical-grade analogue, MRTX-849 (30 mg/kg daily). The results revealed a modest reduction in the rate of growth by single agent KRAS^G12C^ inhibitor (data not shown) and indicates that combination therapy with MRTX-849 will be required to achieve clinically significant anti-tumor responses. In this regard, the protein tyrosine phosphatase, PTPN11/SHP2, has emerged as a promising upstream target as it functions to dephosphorylate multiple proteins leading to sustained KRAS GTP cycling and activation^11,37,38^, and highly specific inhibitors are available. SHP2 also plays a role in negatively regulating JAK-STAT signaling through dephosphorylation of activated STAT proteins^39^ and T cell receptor (TCR) signaling within lymphocytes resulting in decreased T cell effector function^14,15^. Both JAK-STAT signaling, and adaptive immunity and T cell function have been shown to be necessary for oncogene-targeted therapy response^14–17^.

To test the effect of the SHP2 inhibitor RMC-4550 in combination with KRAS^G12C^ inhibition, we plated LLC 46 NRAS KO, CMT-KRAS-G12C.54 and mKRC.1 cell lines in 96-well plates and assessed cell growth in response to a range of concentrations of MRTX-1257 and RMC-4550 alone and in combination. Analysis of the resulting data (see Suppl. Fig. S5 for primary data) with Combenefit^40^ revealed strong *in vitro* synergy with combinations of MRTX-1257 and RMC-4550 in these cell lines (Fig. 2A). Two human KRAS-G12C-positive lung cancer cell lines, Calu1 and H2030, were similarly submitted to growth assays with combinations of MRTX-1257 and RMC-4550. Analysis of the data with Combenefit indicated synergistic drug interactions in these human cell line models as well (Suppl. Fig. S6). Also, combination of these inhibitors yielded greater inhibition of pERK in mKRC.1 and LLC 46 NRAS KO over time compared to either single agent alone (Fig. 2B), suggesting this combination induces synergistic growth, in part, through prevention of ERK reactivation.

**Figure 2.**
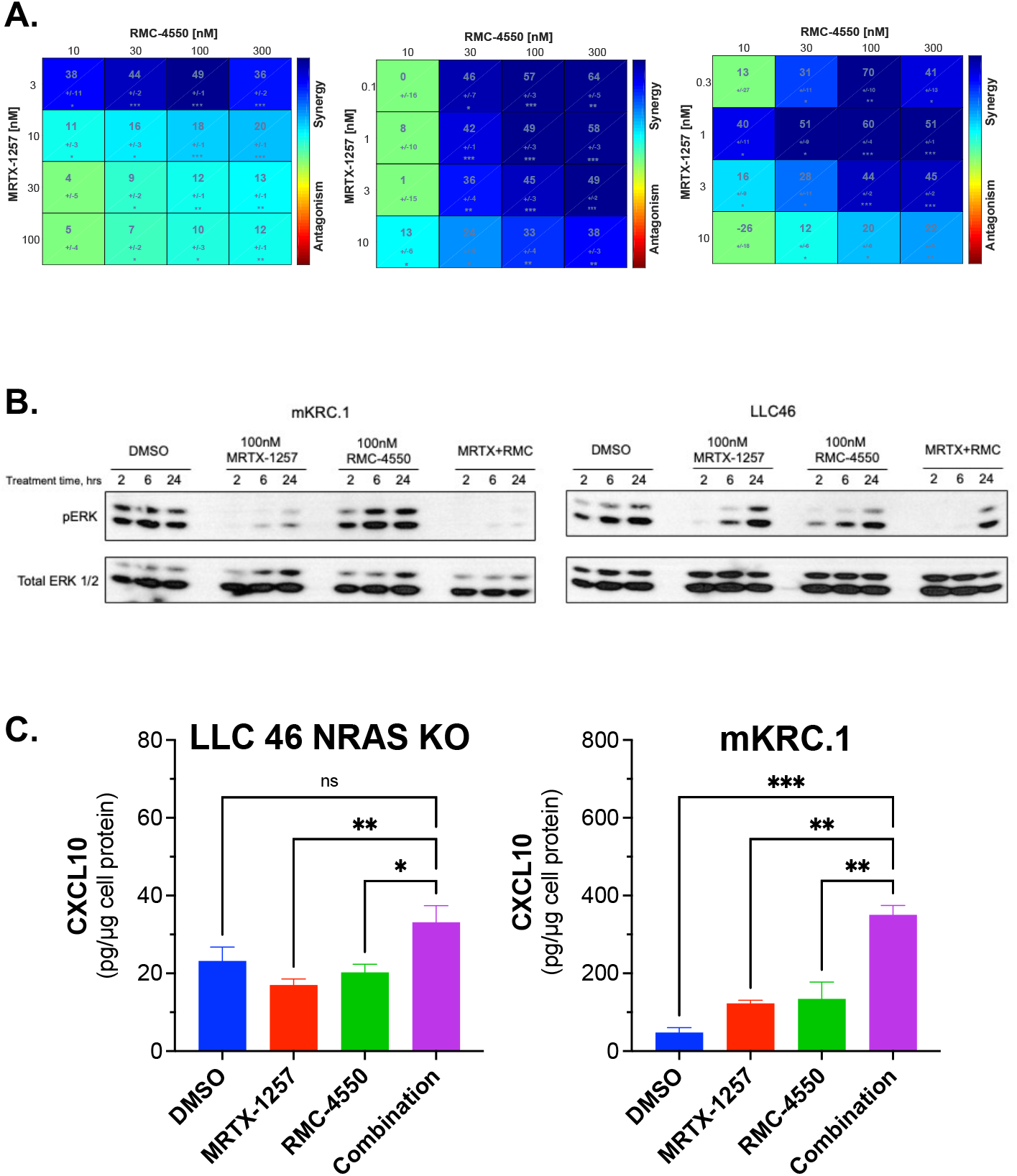
SHP2 inhibition functions synergistically with KRAS-G12C inhibitors in KRAS-G12C-driven NSCLC cell lines and expands chemokine production. (A) Murine cell lines driven by KRAS G12C were plated at 100 cells per well in a 96 well plate and treated with increasing concentrations of either MRTX-1257, RMC-4550, or in combination for 7-10 days, and then assessed for cell growth via CyQuant cell growth assays. HSA synergy was calculated using Combenefit software. (B) LLC 46 NRAS KO and mKRC.1 cells were plated and treated with DMSO control, MRTX-1257 (100nM), RMC-4550 (100nM), or the combination over 2, 6, and 24 hours. Cell lysates were collected and immunoblotted for phospho-ERK and total ERK1/2 protein. (C) LLC 46 NRAS KO and mKRC.1 cells were plated in 6 well plates and treated with either DMSO control, RMC-4550 (100 nM), MRTX-1257 (100 nM), or the combination and allowed to incubate for 2 days. Conditioned cell media was collected and assessed for CXCL10 chemokine expression by ELISA. The data are the mean and SEM of 6 and 3 replicates for LLC 46 NRAS KO and mKRC.1, respectively, and were submitted to 1-way ANOVA and multiple comparisons test.

The combination of MRTX-1257 and RMC-4550 also stimulated significant induction of the anti-tumorigenic chemokine, CXCL10^41^ compared to single agent MRTX-1257 or RMC-4550 in LLC 46 NRAS KO and mKRC.1 (Fig. 2C). It is noteworthy that mKRC.1 cells secrete ~10-fold more CXCL10 compared to LLC 46 NRAS KO. These data show that RMC-4550 in combination with MRTX-1257 results in strong inhibition of cell growth in our models and greater induction of CXCL10 *in vitro*.

### Therapeutic response to KRAS-G12C inhibitor, alone and in combination with SHP2 inhibitor in orthotopic *in vivo* models

Based on the synergistic activity of this combination *in vitro* (Fig. 2A), the drug combination was tested in LLC 46 NRAS KO cells using an orthotopic lung tumor model^29^. Following inoculation of LLC 46 NRAS KO cells into the left lung of C57BL/6 mice, tumors were assessed for growth over time using µCT imaging.

Representative µCT images from this experiment are presented in Supplemental Figure S7. Treatment with single agent RMC-4550 or MRTX-849 resulted in marginally reduced rate of tumor growth compared to diluent control (Fig. 3A). However, the combination of the two agents yielded initial tumor regression and suppressed tumor growth until tumors ultimately progressed (Fig. 3A). Assessment of the individual tumor responses at day 21 (11 days of treatment) among the experimental groups revealed no significant change in tumor volume with single agents compared to diluent control mice, although modest improvement in overall survival was observed (Fig. 3B-C). By contrast, four of the eight mice treated with the combination of MRTX-849 and RMC-4550 experienced tumor shrinkage and significantly improved survival compared to diluent control and each single agent alone (Fig. 3B-C). These data demonstrate that improved MAPK blockade from this combination as well as potentially increased anti-tumor immune cell recruitment through previously outlined mechanisms help drive responses in this murine model of KRAS^G12C^ LUAD.

**Figure 3.**
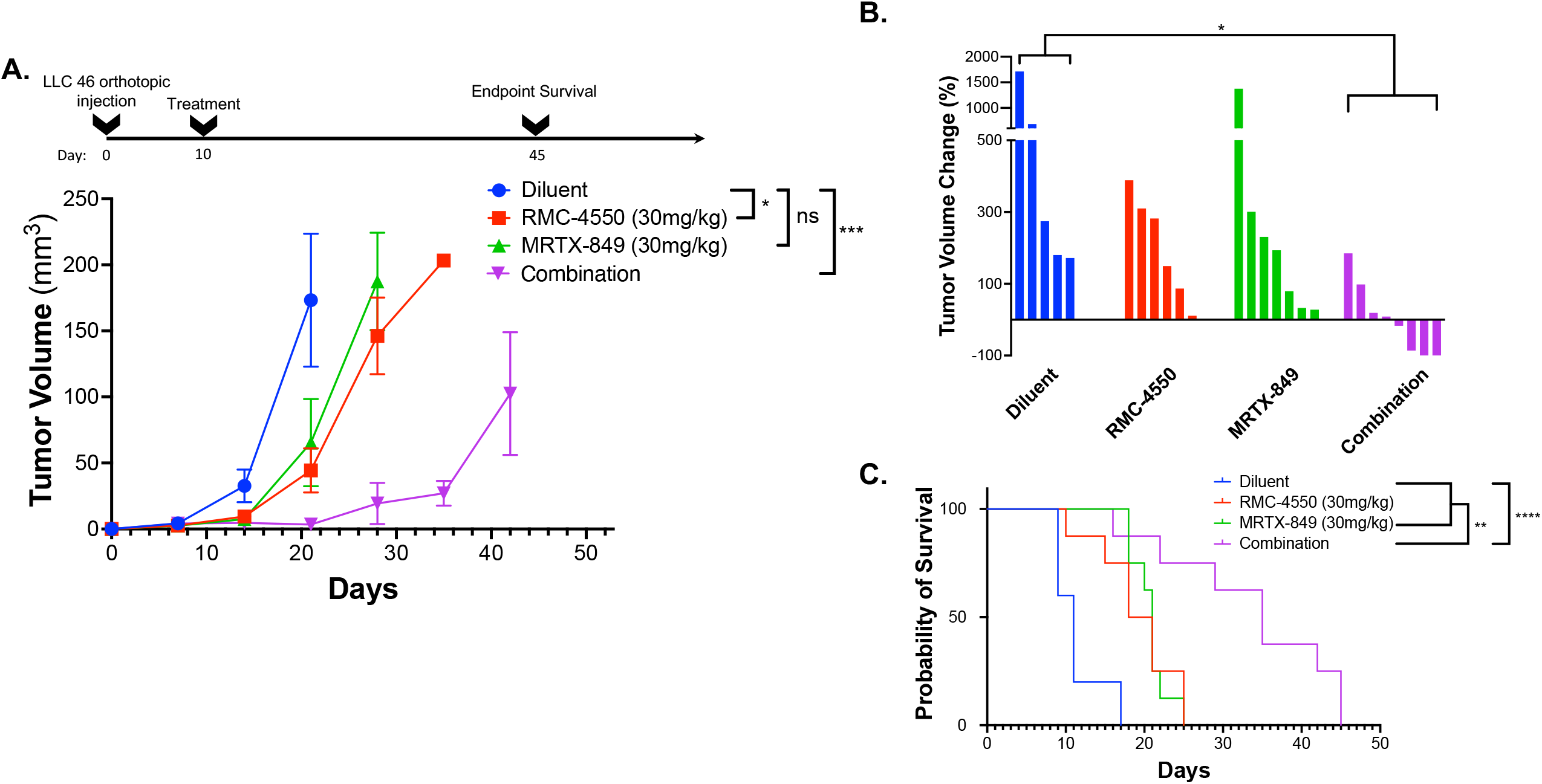
KRAS^G12C^ inhibitors used in combination with SHP2 inhibitors demonstrate improved efficacy compared to either treatment alone *in vivo*. LLC 46 cells were orthotopically implanted into the left lung of C57BL/6 mice and were allowed to establish for 7 days before initial pre-treatment micro-CT imaging. **(A)** Scheme showing the timeline of the experiment overlayed on tumor volume growth curves of Diluent (n=5), RMC-4550 (n=6), MRTX-849 (n=7), and combination-treated (n=8) mouse cohorts. The data were analyzed by 2-way ANOVA with Tukey’s multiple comparisons test. **(B)** Waterfall plot showing precent change in tumor volume from day 21 to baseline (initial pre-treatment size) on day 7. The data were analyzed by Kruskal-Wallis test with Dunn’s multiple comparisons test. **(C)** Survival curves comparing all 4 groups over the course of the study with one-way ANOVA and multiple comparisons test.

In contrast to the LLC 46 NRAS KO model, orthotopic lung tumors established with the mKRC.1 cell line exhibited significant single-agent MRTX-849 responses with all five tumors exhibiting shrinkage after 11 days of treatment (Fig. 4A and B; representative µCT images in Suppl. Fig. S7). Notably, after 7 additional days of MRTX-849 treatment, four of the five tumors exhibited increased volumes, suggesting rapid progression through acquired resistance.

**Figure 4.**
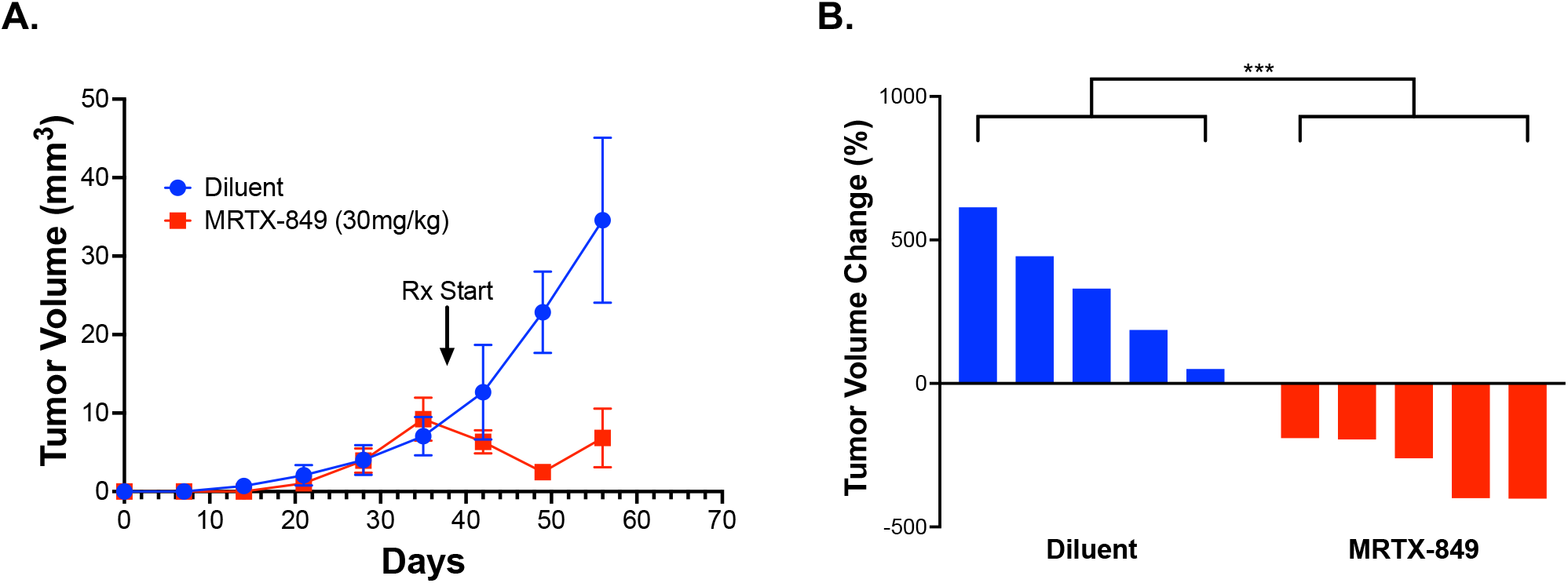
mKRC.1 tumors in C57BL/6 mice show strong response to single agent MRTC-849 treatment. mKRC.1 cells were orthotopically implanted into the left lungs of C57BL/6 mice and allowed to establish for 35 days before initial pre-treatment micro-CT imaging. (**A**) Tumor volumes (mean and SEM) of diluent and MRTX-849 (30mg/kg)-treated mouse cohorts (n=5 per group). (**B**) Waterfall plot showing percent change in tumor volume from day 49 to initial pre-treatment volumes on day 35. The data were analyzed by an un-paired t-test.

### Benefit from SHP2i and KRAS^G12C^ inhibitor combinations requires adaptive immunity

To test the role of adaptive immunity in driving tumor response to combined KRAS^G12C^ and SHP2 inhibition, LLC 46 NRAS KO cells were inoculated into the left lung of athymic *nu/nu* mice and tumors were allowed to establish. Subsequently, tumor-bearing nu/nu mice were treated with diluent control, RMC-4550, MRTX-849, or the combination. While either drug alone modestly reduced tumor growth similar to the result in C57BL/6 mice (Fig. 3A), the combination failed to yield any additional benefit (Fig. 5A). This was further supported by assessing percent tumor volume change in the individual mice, as the combination treatment group was not different than the individual tumors within the single agent groups (Fig. 5B). These results indicate that adaptive immunity is necessary for the enhanced response to this combination observed in immune competent hosts.

**Figure 5.**
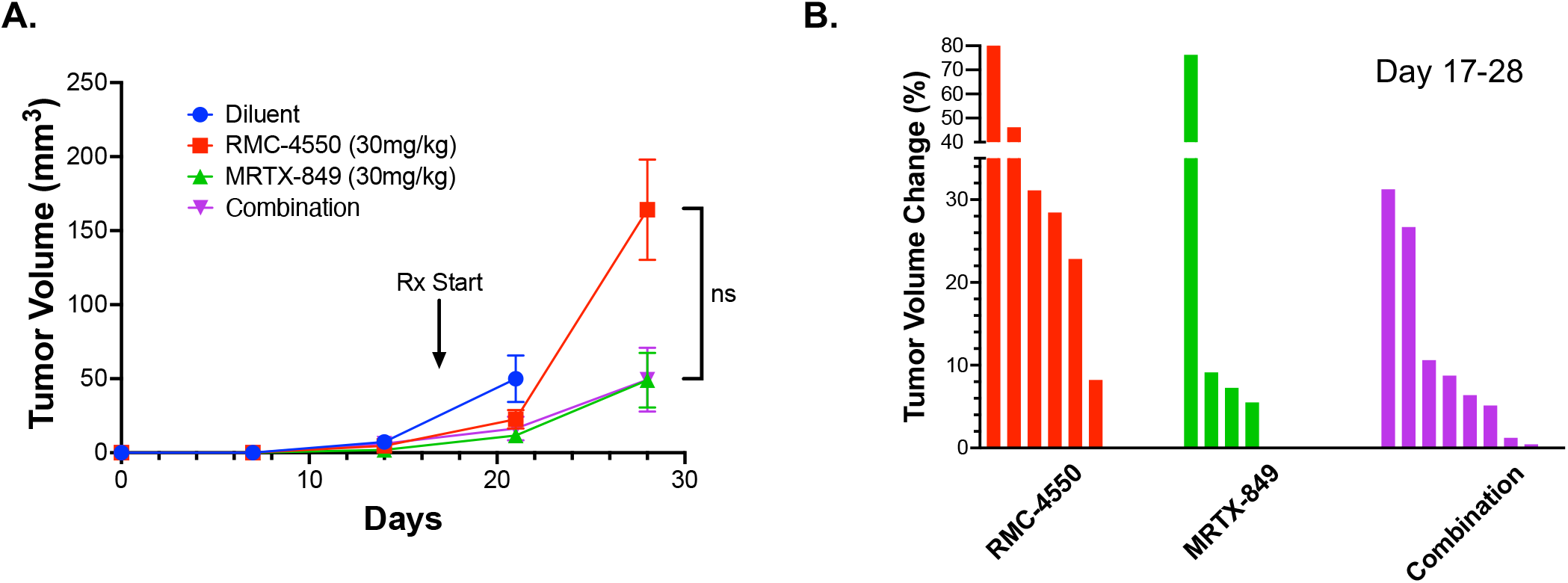
LLC 46 NRAS KO tumors in nude mice demonstrate decreased response to MRTX-849 and/or RMC-4550. LLC 46 cells were orthotopically implanted into the left lung *of nu/nu* mice and allowed to establish for 7 days before initial pre-treatment micro-CT imaging. (**A**) Tumor volumes (mean and SEM) of diluent (n=8), RMC-4550 (30mg/kg; n=8), MRTX-849 (30mg/kg; n=10), and combination-treated (n=8) mouse cohorts. (**B**) Waterfall plot showing percent change in tumor volume from initial pre-treatment on day 17 to treatment day 28.

## Discussion

The murine KRAS^G12C^-driven lung cancer cell lines developed in this study provide novel models to investigate KRAS^G12C^ inhibitor responsiveness in the immune competent setting. This is critical since oncogene-targeted agents including KRAS^G12C^ inhibitors have been shown to initiate functional interactions with host immunity^26,27,33^. The new lung cancer cell lines described herein uniformly exhibit high *in vitro* sensitivity to KRAS^G12C^ inhibitors and altered growth signaling in CMT KRAS^G12C^ clones compared to parental G12V cells. This is most likely due to varied downstream signaling dynamics among G12 mutations^8^ as well as disruption of the unedited KRAS-G12V allele as a result of CRISPR-Cas9 editing yielding a single functional KRAS-G12C allele. Similar to observed clinical activity of single agent KRAS^G12C^ inhibitors in which overall response rates are lower than 45%, and the durability of treatment response is rather brief, our murine models replicate these differential responses through initial progressive disease (LLC NRAS KO) and rapid progression after initial shrinkage (mKRC.1) to MRTX-849 as a single agent. Thus, these new murine KRAS^G12C^-driven lung cancer cell lines propagated as orthotopic tumors reflect treatment responses in KRAS^G12C^-positive patients treated with single agent KRAS^G12C^ inhibitors ^5,42^. The Downward group previously deleted NRAS in a derivative cell line of the murine LLC model to generate a KRAS^G12C^ inhibitor-sensitive cell line^27,33^. Similar to our findings with LLC 46 NRAS KO cells, their LLC derivative exhibited modest *in vivo* sensitivity to KRAS^G12C^ inhibitor monotherapy. Also, KRAS mutant murine CT26 cells of colorectal origin have been engineered to express the KRAS^G12C^ allele and exhibited single agent KRAS^G12C^ inhibitor activity in syngeneic mice^26^. Two distinct KRAS^G12C^ GEMMs have been published^34,43^, although the mKRC.1 cell line described herein appears to be the sole stable cell line derived from a GEMM. In summary, the novel murine KRAS^G12C^ cell lines described herein add to the collection of models to further explore experimental therapeutics involving KRAS^G12C^ inhibitors.

The studies with the LLC NRAS KO cells propagated orthotopically in C57BL/6 mice demonstrated that combined MRTX-849 and SHP inhibitor, RMC-4550, induced tumor shrinkage, albeit transiently (Fig. 3). Notably, this synergistic response was not observed when the experiment was performed in athymic *nu/nu* mice (Fig. 5), indicating a role for adaptive immunity for full therapeutic efficacy. In fact, a role for innate and adaptive immunity in contributing to anti-tumor responses to oncogene-targeted therapeutics is documented in the literature^16-22,26,27^. In addition, our data show that KRAS^G12C^ inhibitors variably induce chemokines (CXCL10) capable of recruiting anti-tumorigenic immune cells such as NK, CD8 T, and dendritic cells into the TME, which is exacerbated in combination with SHP2 inhibitor (Fig. 2C). Further, mKRC.1 cells more strongly induced CXCL10 upon i*n vitro* MRTX-1257 and MRTX-1257/RMC-4550 combination treatment and may account for the single agent MRTX-849 activity *in vivo* compared to LLC NRAS KO cells. Thus, the extent of chemokine induction following KRAS G12Ci may contribute to the depth and durability of response. Canon et al. demonstrated the involvement of anti-tumor immunity in the therapeutic response to sotorasib (AMG-510) using the CT-26 cell line model^26^. Also, our own studies with murine EML4-ALK-driven cell lines demonstrate a requirement for adaptive immunity for durable responses to the ALK inhibitor, alectinib^19^. Current understanding surrounding the depth and duration of response to oncogene-targeted inhibitors suggests that the extent to which chemokine induction occurs with oncogene-specific therapies associates with the degree of response, as EGFR mutant lung cancer patients exhibiting greater interferon γ transcriptional responses to TKIs presented with longer progression-free survival. Identifying why some patients experience stronger immunogenic responses to oncogene-targeted inhibitors remains an unanswered question in the field and a deeper understanding may unveil novel mechanisms to increase objective responses and durability of treatment with oncogene-targeted agents in general.

Precision therapy with oncogene-targeted drugs has dramatically changed the outcomes for lung cancer patients with tumors positive for mutated EGFR or fusion kinases including ALK and ROS1 and others. In fact, TKIs are now FDA-approved first-line therapies for these subsets of lung cancer. By contrast, KRAS^G12C^ inhibitors remain second-line treatments for KRAS^G12C^-positive lung cancers due to lower objective response rates and shorter durations of treatment response relative to the clinical activity observed with anti-PD1-based immunotherapy, either alone or in combination with cytotoxic chemotherapy^5^. Completed clinical trials of sotorasib and adagrasib reveal response rates ranging from 37 to 43% and rather brief PFS (6 – 7 months) relative to that associated with TKIs and RTK-driven lung cancers^5^. The lower efficacy of sotorasib and adagrasib as monotherapies supports their combination with other pathway-targeted agents or immune therapy for better anti-cancer activity. Inhibition of the PD-1/PD-L1 axis remains a primary treatment strategy for KRAS mutant lung cancer. Numerous studies have shown that the existence of an active IFNY signature within the TME is required for response to immune checkpoint inhibitors^44,45^. Because oncogene-targeted inhibitors have been shown to induce an IFNY signature with the TME^16,17^, numerous trials combining immune therapies with KRAS^G12C^ inhibitors have been initiated to see if KRAS inhibition can expand checkpoint inhibitor responses.

Among the targets being investigated for combined benefit with KRAS-G12Ci’s, SHP2 inhibitors have gained interest due to their ability to impact tumor cell autonomous MAPK signaling and JAK-STAT signaling, as well as regulation of T cell receptor signaling^6,9,14,15,20,46,47^. Multiple SHP2 inhibitors are now in the clinical trial pipeline being tested as monotherapies, and in combination with KRAS^G12C^ inhibitors^47^. In addition, emerging data supports the rationale of targeting upstream of KRAS GTP nucleotide cycling as KRAS^G12C^ inhibitors covalently bind KRAS in its GDP-bound (inactive) form^48^. Unlike other codon 12 missense mutations, KRAS^G12C^ maintains most of its intrinsic GTP hydrolysis activity, allowing the KRAS protein to inactivate through GTP hydrolysis even in its mutant form^8^. This represents an interesting therapeutic vulnerability as upstream blockade of KRAS signaling should impede KRAS GTP nucleotide cycling into its active form resulting in a higher ratio of KRAS-GDP – the substrate for KRAS^G12C^ inhibitors. In this regard, the SHP2 inhibitor RMC-4550 exhibits strongly synergistic in vitro growth inhibition in combination with MRTX-1257 in all three cell lines tested. Moreover, RMC-4550 markedly increases efficacy of MRTX-849 in the LLC NRAS KO orthotopic C57BL/6 model (Fig. 3A), but not immune deficient hosts (Fig. 5A), demonstrating the requirement of host immunity for this therapeutic effect. The pre-clinical KRAS^G12C^ models described in this report allow for deeper experimental exploration of the impacts of inhibiting SHP2 on both tumor cell autonomous signaling and non-autonomous targets such as T cell signaling. These models may also serve as pre-clinical models for other KRAS^G12C^ inhibitor combinations currently in the clinical trial pipeline such as inhibitors of EGFR and SOS1^10^.

## Author Contributions

Conception and design: D.J.S., T.K.H, E.K.K., R.A.N, L.E.H

Development of methodology: D.J.S., T.K.H, A.T.L., R.A.N.,

L.E.H Acquisition of data: D.J.S., T.K.H, L.E.H

Analysis and interpretation of data: D.J.S., T.K.H, L.E.H

Writing, review, and/or revision of the manuscript: D.J.S., T.K.H, E.K.K., R.A.N, L.E.H

## Acknowledgements

The RNAseq was performed by the University of Colorado Cancer Center Genomics shared resource.

## Data Availability

RNA-seq data have been deposited in the Gene Expression Omnibus database (GSE207098). All other data and cell lines supporting findings of the study are available within the article, in the supplemental information files, or from the corresponding author upon request.

