## Supplementary Figures for "Evaluation of KRAS^G12C^ Inhibitor Responses in Novel Murine KRAS^G12C^ Lung Cancer Cell Line Models"

Daniel J. Sisler

Department of Craniofacial Biology

University of Colorado Anschutz Medical Campus

Aurora, Colorado

(303) 724-4632

**Running Title:** Novel murine KRAS^G12C^-dependent lung cancer cell lines

**Supplemental Figures S1 – S7**

**Supplemental Table S1**

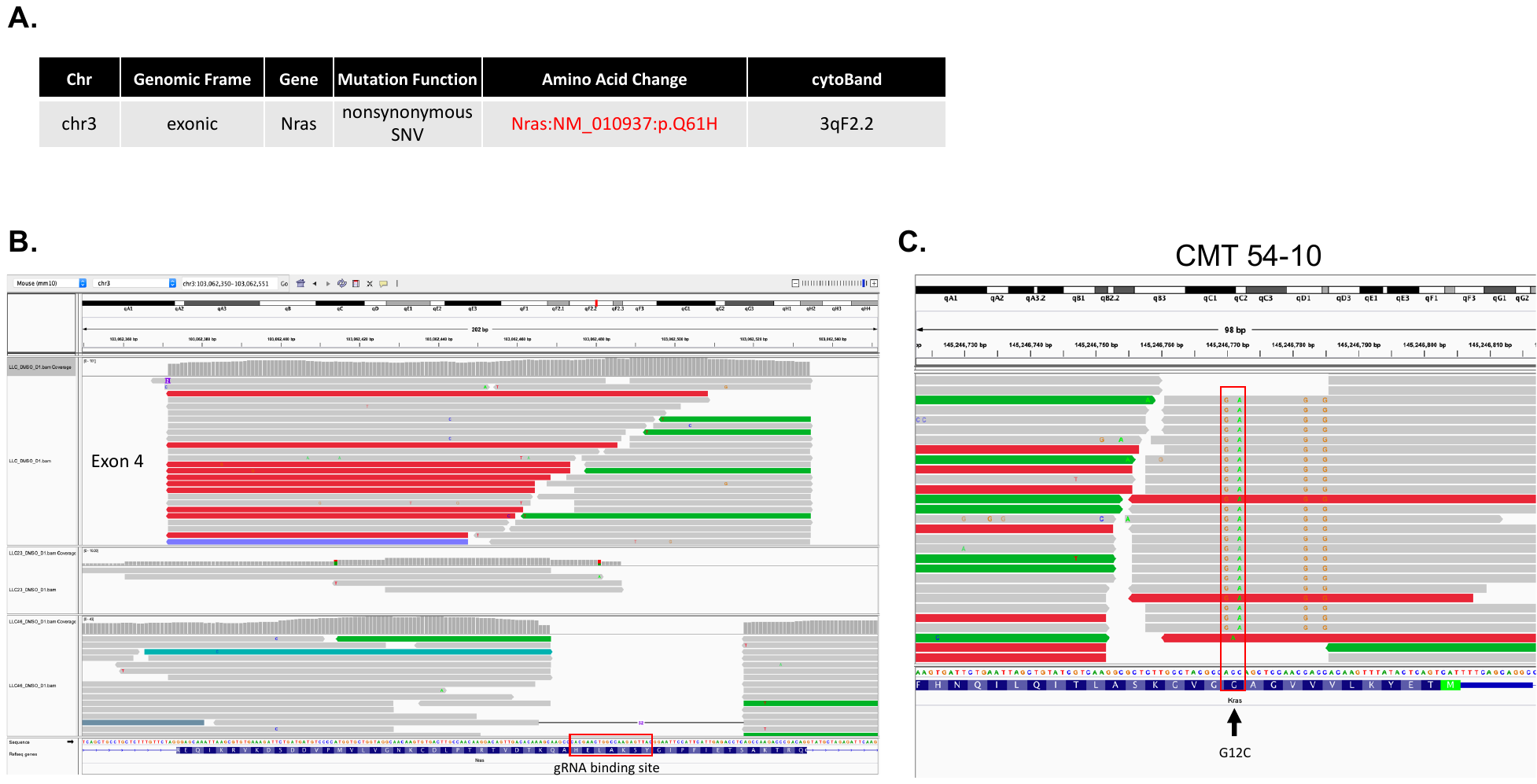

**Supplemental Figure S1. Validation of NRAS Q61H mutation and CRISPR/Cas9 genomic editing.** (**A**) Variant call analysis (See Materials and Methods) was performed on parental LLC RNA sequencing data. (**B**) Integrated genomic viewer (IGV) analysis of NRAS exon 5 in parental LLCs compared to 2 LLC NRAS KO subclones (LLC 23 and LLC 46). (**C**) IGV analysis of CMT KRAS G12C subclone 54-10 RNA sequencing data focused on KRAS exon 2, codon 12.

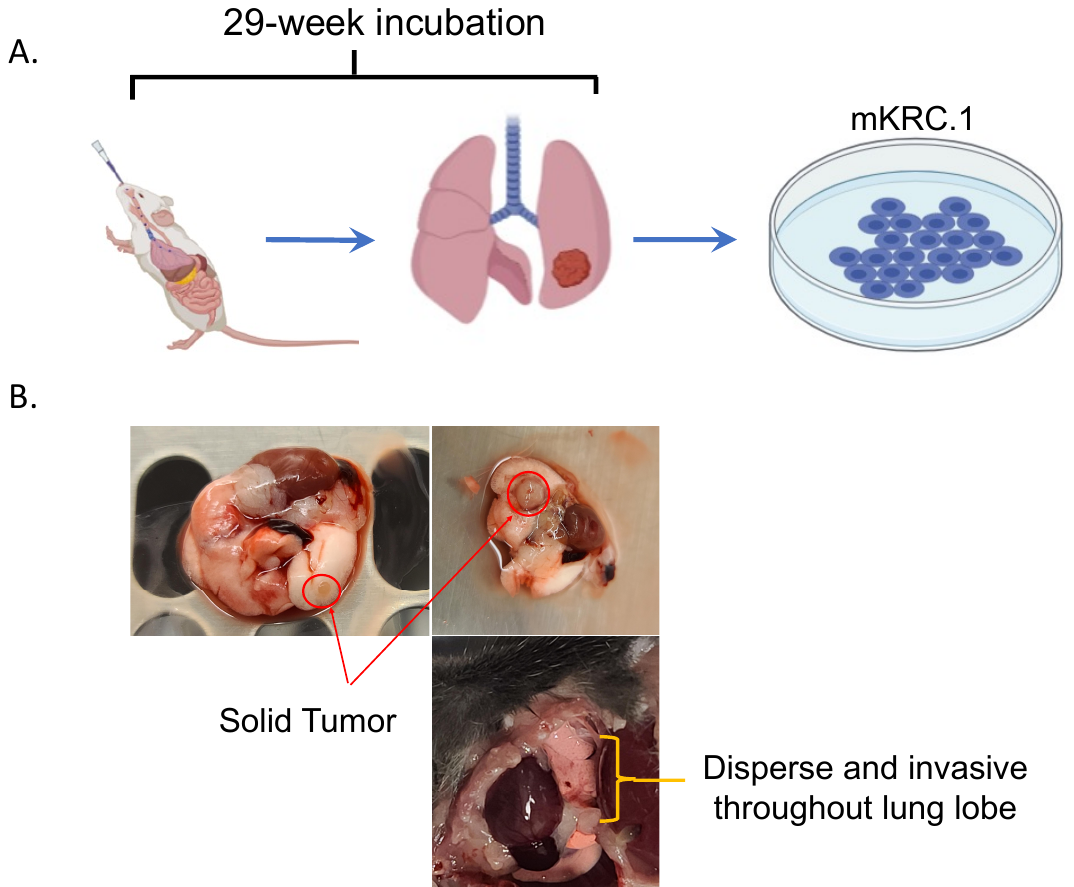

**Supplemental Figure S2. Overview of development of mKRC.1 cell line.** (**A**) Schematic of intratracheal injections followed by lung resection and cell line development. (**B**) Gross images showing lung tumor nodules and patterns of tumor development within mouse lungs.

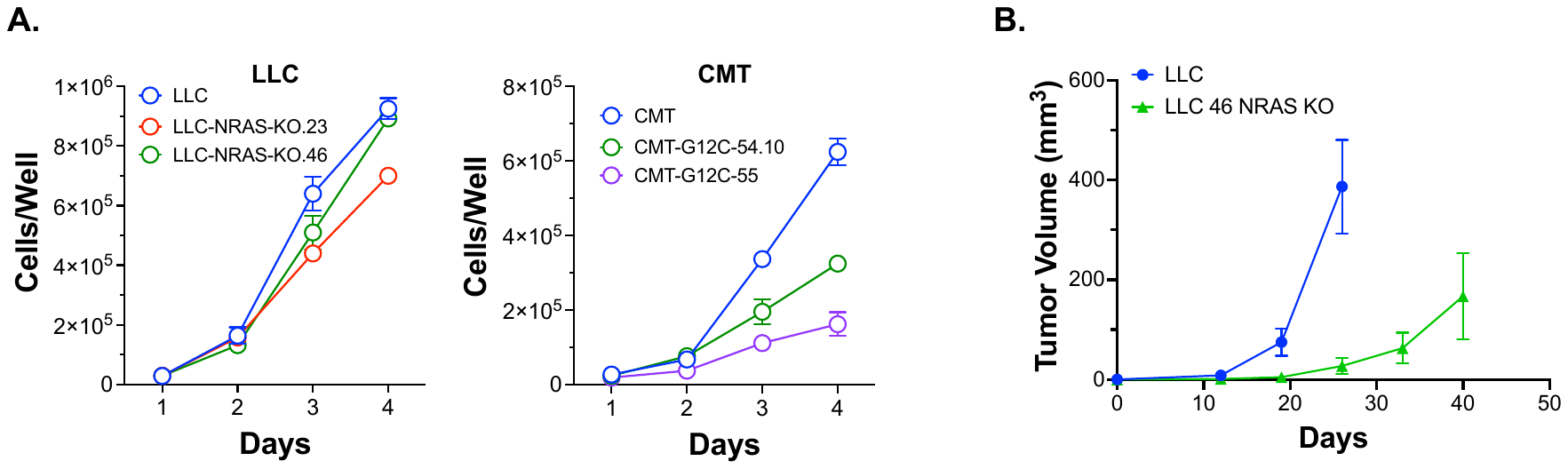

**Supplementary Figure S3. Baseline growth rates of LLC and CMT167 CRISPR-edited cell lines *in vitro* and *in vivo*.** (**A**) Parental and CRISPR-edited cell lines were plated at 25,000 cells per well in 24-well plates and cultured. Duplicate wells were trypsinized and cells counted after 1-4 days. The data are the means and SEM. (**B**) LLC parental and LLC 46 NRAS KO cell lines were orthotopically implanted into the left lungs of C57BL/6 mice and tumor growth was measured via weekly μCT imaging. The data are the means and SEM of 5 mice per cell line.

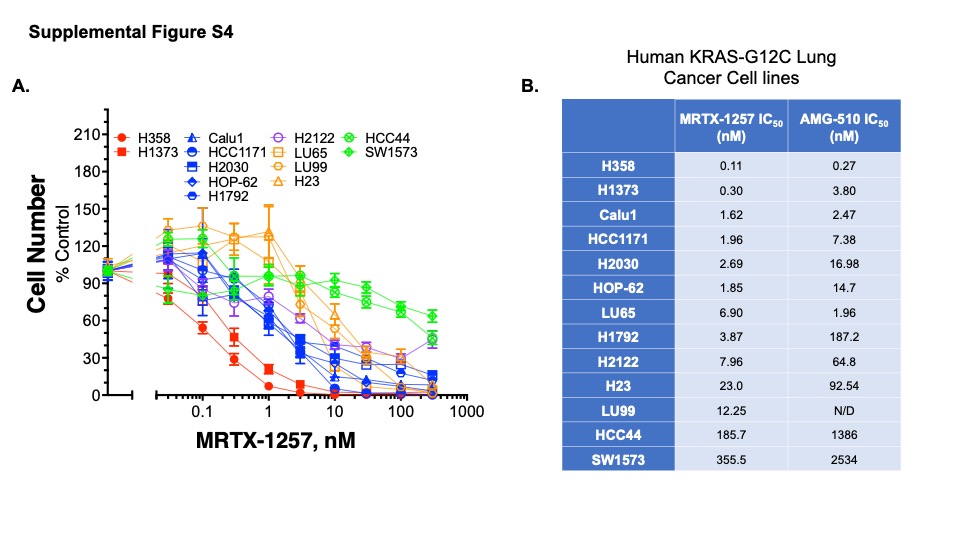

**Supplemental Figure S4. Sensitivity of human KRAS-G12C positive lung cancer cell lines to MRTX-1257 and AMG-510.** A panel of thirteen human KRAS-G12C mutant lung cancer cell lines were seeded at 100-200 cells/well in 96-well plates and treated for 7-10 days with MRTX-1257 (**A**) or AMG-510 (not shown). Cell number was assayed with CyQUANT reagent. The data are the means and SEM of triplicate measurements presented as percent of the DMSO control treatments. (**B**) Prism 9 was used to calculate the IC_50_ values from the MRTX-1257 and AMG-510 dose-response curves.

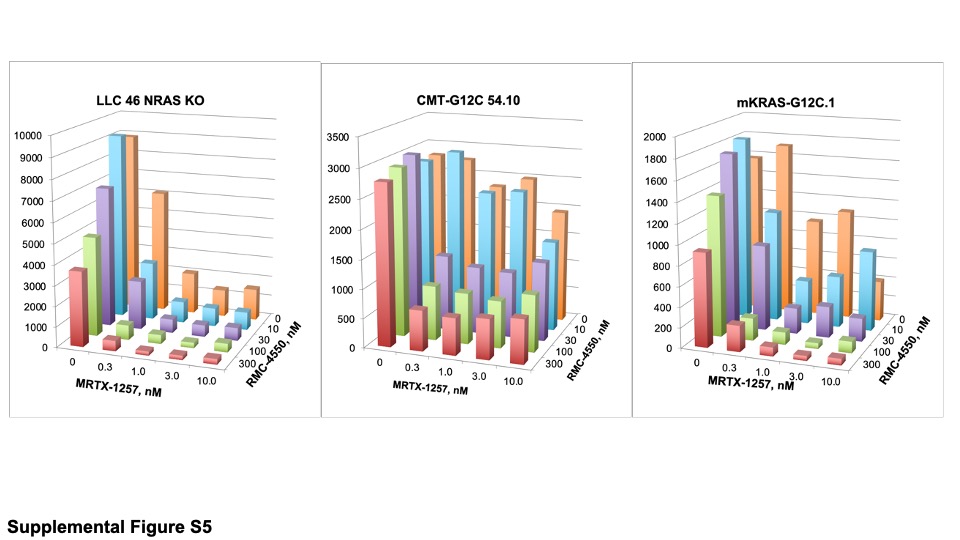

**Supplemental Figure S5. Primary MRTX-1257-RMC-4550 combination data used for calculating drug synergy.** The indicated murine KRAS-G12C cell lines were treated with combinations of MRTX-1257 and RMC-4550 at the concentrations shown in quadruplicate in a 96-well format. After 7-10 days of treatment, cell number was measured with CyQUANT reagent and the average values among the replicates is plotted as shown.

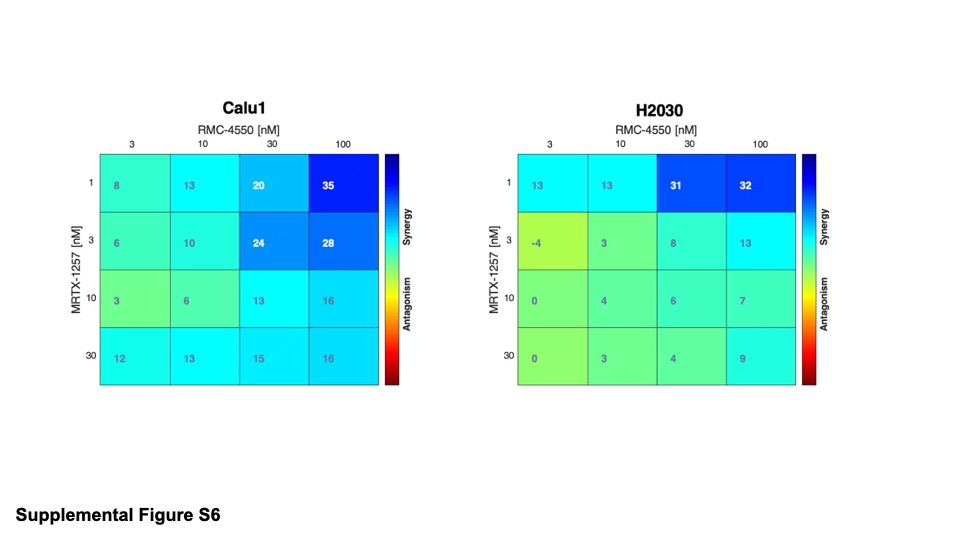

**Supplemental Figure S6. Analysis of KRAS-G12C-SHP2 inhibitor synergy in human KRAS-G12C lung cancer cell lines.** Calu1 and H2030 were seeded at 100 cells/well in 96-well plates and treated in duplicate with the indicated concentrations of MRTX-1257 and RMC-4550 for 7-10 days. Cell numbers were assayed with CyQUANT reagent and the resulting data were analyzed with Combenefit for drug synergy using the HSA model.

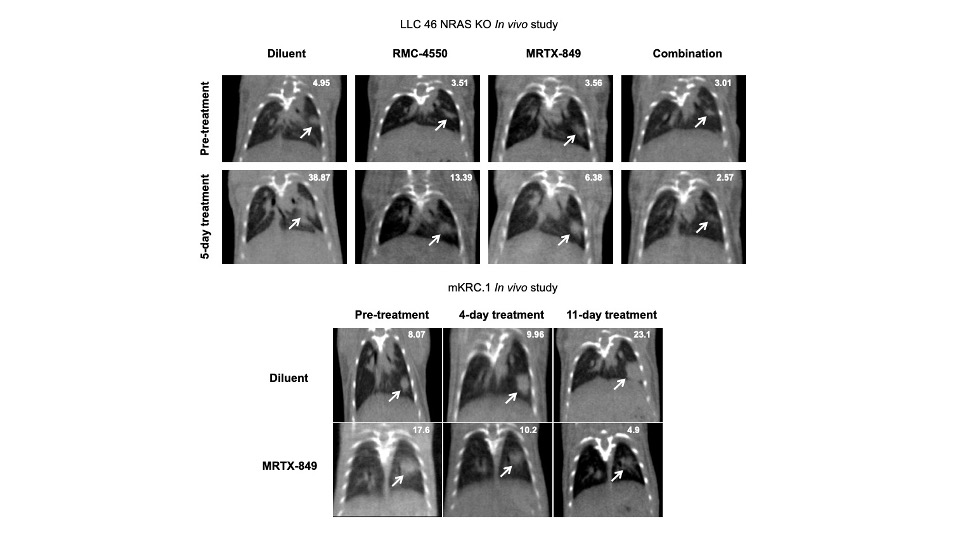

**Supplemental Figure S7. MicroCT images from representative orthotopic LLC 46 NRAS KO and mKRC.1 lung tumors.** Representative orthotopic lung tumors from the LLC 46 NRAS KO and mKRC.1 experiments in Figures 3 and 4, respectively are shown. The top panel shows representative tumor-bearing mice from the LLC 46 NRAS KO experiment from the four experimental groups before and after 5 days of daily treatment. The lower panel shows μCT images representative of diluent and MRTX-849 treated mice prior to and after 4 and 11 days of treatment. The values in the upper right-hand corner of each image are the tumor volumes calculated by the ITK-SNAP software program.

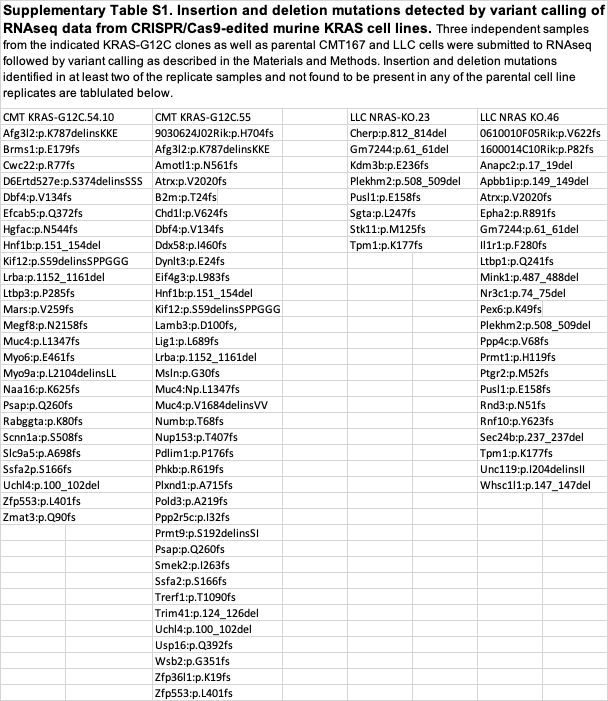
